# Maternal arterial blood values during delivery: effect of mode of delivery, maternal characteristics, obstetric interventions and correlation to fetal umbilical cord blood

**DOI:** 10.1101/2020.02.14.950089

**Authors:** Mehreen Zaigham, Sara Helfer, Karl Heby Kristensen, Per-Erik Isberg, Nana Wiberg

## Abstract

**Objective:** To determine a reference interval for maternal arterial blood values during vaginal delivery and to elucidate the effect of common maternal characteristics and obstetric interventions on maternal acid base values during vaginal and planned cesarean section (CS).

**Design:** Prospective, observational study of randomly selected women undergoing vaginal deliveries and planned CS at Skåne University Hospital, Malmö, Sweden.

**Results:** Two hundred and fifty women undergoing vaginal delivery (VD) and fifty-eight women undergoing planned CS were recruited. We found significant differences for gestational age, parity, artery pH, pCO_2_, pO_2_, sO_2_ and cord venous pH, pCO_2_ and lactate between the two study groups (*P* < 0.005). For women undergoing vaginal delivery, we found significant changes in base deficit, hemoglobin, bilirubin, potassium, glucose and lactate values as compared to women with planned CS (*P* < 0.02). Maternal characteristics did not significantly affect acid base parameters however, multiple regression showed significant associations for the use of epidural anesthesia on maternal pH (*P* < 0.05) and pO_2_ (*P* < 0.01); and synthetic oxytocin on pCO_2_ (*P* = 0.08), glucose (*P* < 0.00) and lactate (*P* < 0.02) in maternal blood. Maternal arterial pH, pCO_2_ and lactate correlated significantly to values in venous umbilical cord blood (*P* < 0.000).

**Conclusions:** Reference values for maternal arterial blood gases in vaginal deliveries for term pregnancies were outlined and we found that most arterial blood gas parameters varied significantly according to mode of delivery. The use of different obstetrical interventions like epidural anesthesia or synthetic oxytocin, resulted in significant changes in blood gas values.

## Introduction

It is well known that there are significant changes in almost all maternal organ systems during pregnancy and that these physiological changes enable the mother to optimally nourish the fetus as well as prepare her for labor (1). Klajnbard and Maguire (2, 3) showed large variations in most maternal venous parameters during the three trimesters of pregnancy. Despite this knowledge, obstetricians routinely use reference values from non-pregnant women to assess the condition of the pregnant patient. Few studies have focused on physiological variations during pregnancy and delivery, and arterial blood gases including electrolytes, bilirubin, glucose and lactate have seldom been studied. In addition, the possible impact of mode of delivery on different maternal and obstetric factors commonly encountered during delivery, has not been studied.

One of the most frequently encountered maternal risk factors during pregnancy and delivery is obesity. Maternal obesity is associated with increased fat deposition, lower body muscle mass, decreased respiratory capacity and chronic inflammatory changes with an increased risk for prolonged labor and adverse maternal-neonatal outcome (4). Smoking during pregnancy is another potentially detrimental risk factor which can affect maternal acid base balance via vasoconstrictive effects on blood vessels and it is well recognized that maternal smoking is associated with hypertension and decreased lung capacity (5, 6).

Among obstetric factors, synthetic oxytocin used to augment labor, has powerful effects on uterine contractions and maternal physiology (7). Similarly, the use of spinal/epidural anesthesia (EDA) to provide intrapartum analgesia can result in decreased peripheral vascular resistance which in turn causes alterations to uteroplacental blood flow with impaired oxygenation (8). EDA is well known to prolong labor and increase the risk for adverse maternal outcome and obstetric intervention (7, 9).

Thus, the main aim of this study was to determine reference values for maternal arterial blood values during vaginal delivery. We also wanted to elucidate the effect of common maternal characteristics and obstetric interventions on maternal acid base values during vaginal and planned CS. Correlations between maternal acid base values and cord venous blood were also explored.

## Materials and Methods

### Study population

The study was carried out at the Department of Obstetrics and Gynecology, Skåne University Hospital, Sweden. Our study population consisted of two subgroups: patients intended for vaginal delivery (VD) from whom the reference interval was calculated (February 2010 to September 2011) and patients undergoing planned caesarean section (CS) (October 2006 to January 2007 and December 2009 to July 2011). It was established from a previously published paper that the demographic and obstetrical data of the VD subgroup were comparable to the background population i.e. non-participating women who gave birth at the hospital during the same time period as the study (10).

Swedish speaking women in active labor with a cervix dilation of 5-6 centimeters (cm) intended for vaginal delivery alternatively, women intended for planned CS, were randomly informed about the study and participating women were required to give written consent before enrollment in the study. All participants had singleton pregnancies dated by an early second trimester ultrasound, cephalic presentation and non-pathological cardiotocography (CTG) at inclusion.

Immediately after delivery of the baby, before the first cry, a blood gas sample was obtained from the mother’s right radial artery by N.W. Simultaneously, a midwife/junior nurse collected blood from the umbilical cord artery and vein using 2 mL pre-heparinized syringes as per routine at the department. In the planned CS subgroup, a maternal arterial blood gas sample was collected by the anesthesiologist on call at the exact time point when the fetus was delivered with simultaneous sampling from the umbilical cord by N.W. All blood samples were analyzed within 10 minutes after delivery to ensure optimal analysis quality.

For both groups, neonatal and obstetric data of significance were entered in to the study’s database directly after the delivery of the infant whilst the mother was still admitted to the department.

### Biochemical analyses

Maternal and fetal umbilical cord blood were analyzed using the blood gas analyzer, ABL 800 Flex, Radiometer, Copenhagen, Denmark. In Lund, the blood gas analyzer programmed to automatically calculate base deficit in extracellular fluid (BD_ecf_) whilst in Malmö, the ABL was set to calculate base deficit (BD) in whole blood. Since the latter is known to introduce a serious confounding factor (11, 12), we standardized BD values to BD_ecf_ post hoc using the algorithms in the ABL 800 manual:

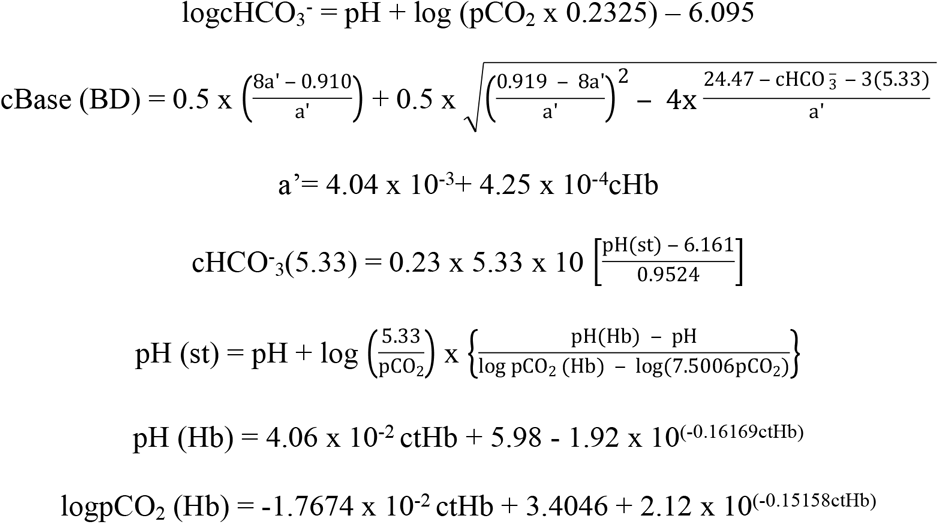

where “c” denotes concentration and “ctHb” denotes total concentration of hemoglobin

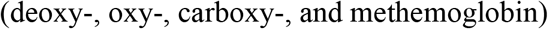

### Statistical analyses

Fisher’s exact test was used for comparison of categorical variables. Group comparison of continuous variables was performed using the Kruskal-Wallis test or the Mann-Whitney *U* test, when appropriate. Values were reported as mean with standard deviation (SD) and median with interquartile range (IQR). Association between variables was reported using regression analysis. Correlation between variables was calculated by Spearman’s test. A two-sided *P*-value < 0.05 was considered significant. Analyses were performed using IBM SPSS Statistics for Windows, version 25.0 (SPSS Inc. Chicago, Illinois, USA).

## Ethical Approval

The study was approved by the Central Ethical Board, Stockholm, Dnr: Ö50-2005.

## Results

A total of 309 women agreed to participate in the study. In the group with planned VD, one case was excluded due to emergency CS performed in general anesthesia, leaving 250 cases for final analysis in the VD group and 58 cases in the planned CS group.

The maternal and fetal characteristics are reported in Table 1.

**Table 1.**
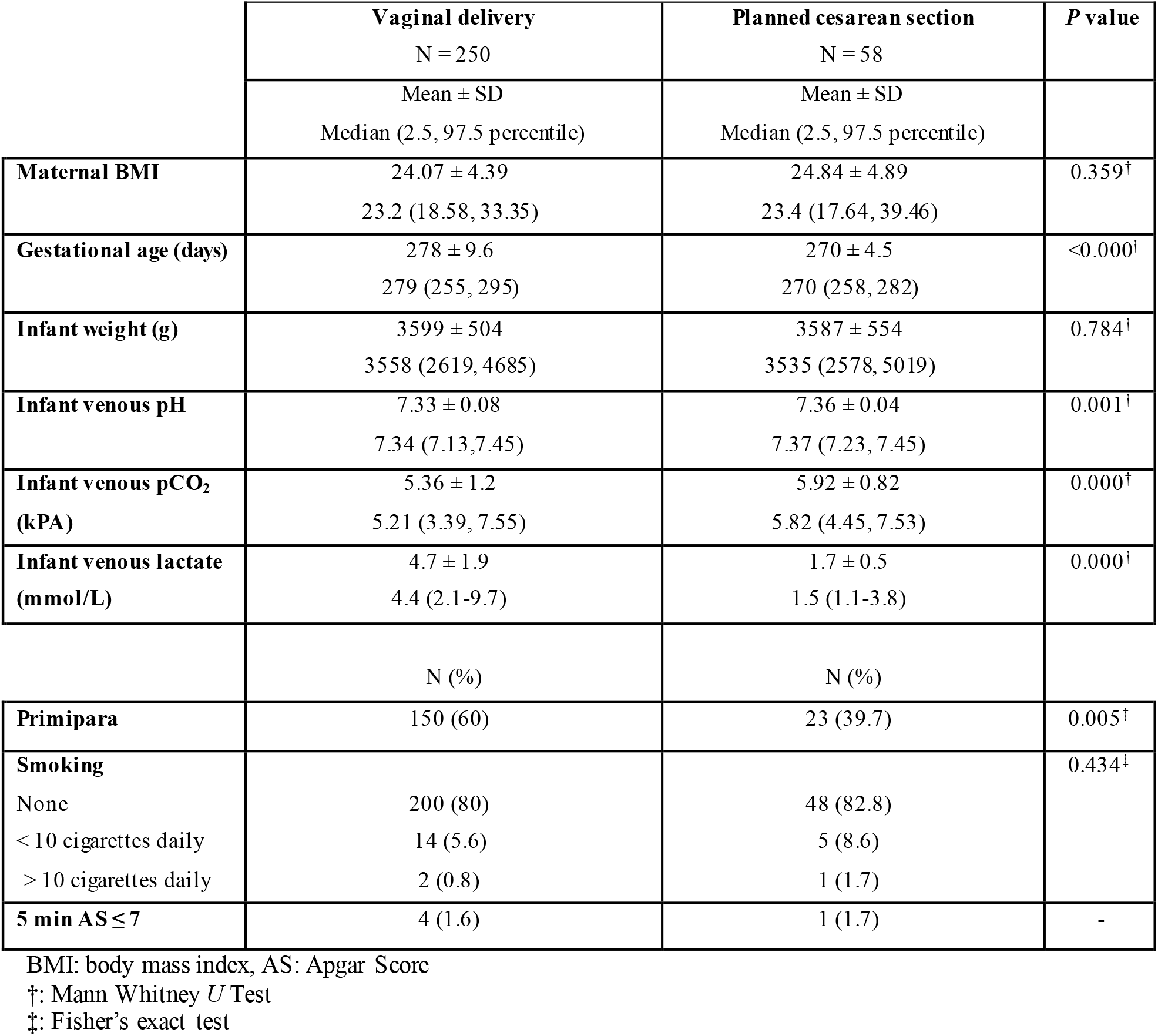
Maternal and fetal characteristics of the two study groups.

Significant differences were seen in the frequency of nulli- versus primipara *(P* < 0.005), gestational age (*P* < 0.000) and for values in umbilical cord venous blood (*P* < 0.008). Although the differences in pH and pCO_2_ were significant, the most remarkable difference was seen for lactate (*P* < 0.000).

Reference values for common maternal acid base values are reported in Table 2.

**Table 2.** Reference values for maternal arterial blood gases according to mode of delivery. Blood collected from radial artery and analyzed by ABL800^™^ (Radiometer, Copenhagen, Denmark). Mann Whitney *U* Test used for two group comparison.

|  | Vaginal delivery<br>N = 250 | Planned cesarean section<br>N = 59 | P value |
| --- | --- | --- | --- |
| | Mean ± SD<br>Median (2.5, 97.5 percentile)<br>[Range] | Mean ± SD<br>Median (2.5, 97.5 percentile)<br>[Range] | |
| **pH** | 7.41 ± 0.06<br>7.41 (7.32, 7.54)<br>[7.32-7.83] | 7.44 ± 0.05<br>7.44 (7.31, 7.57)<br>[7.29-7.57] | < 0.000 |
| **pCO<sub>2</sub> (kPa)** | 3.29 ± 0.52<br>3.35 (2.18, 4.22)<br>[1.03-4.37] | 4.16 ± 0.62<br>4.25 (2.81, 5.95)<br>[2.59-6.45] | < 0.000 |
| **pO<sub>2</sub> (kPa)** | 15.7 ± 4<br>15.6 (11.5, 21.3)<br>[7.3-64.5] | 20.7 ± 7.4<br>19.2 (14.4, 53.7)<br>[14.3-57.5] | < 0.000 |
| **sO<sub>2</sub> (%)** | 98.2 ± 1.1<br>98.4 (96.2, 99.4)<br>[89.4-99.5] | 99.1 ± 0.5<br>99.1 (96.2, 99.4)<br>[97.5-100] | < 0.000 |
| **BD (mmol/L)** | 7.2 ± 2.9<br>7 (3 -13.4)<br>[-10.6, 21.6] | 2.2 ± 1.4<br>2.1 (-1.1, -5.3)<br>[-1.3, 5.6] | < 0.000 |
| **ctHb (g/L)** | 136 ± 10<br>137 (117, 156)<br>[97-190] | 113 ± 19<br>110 (81, 182)<br>[78-196] | < 0.000 |
| **Hctc** | 0.417 ± 0.032<br>0.42 (0.36, 0.48)<br>[0.3-0.55] | 0.35 ± 0.056<br>0.34 (0.25-0.56)<br>[0.24-0.6] | < 0.000 |
| **FHbF (%)** | 6.4 ± 6.3<br>5 (0, 28.4)<br>[0-33] | 7.2 ± 5.3<br>5.5 (0, 18.9)<br>[0-19] | 0.175 |
| **ctBil (μmol/l)** | 13 ± 8<br>12 (0, 35)<br>[0-48] | 10 ± 7<br>9 (0, 34)<br>[0-38] | 0.004 |
| **Na<sup>+</sup> (mmol/L)** | 135 ± 2.6<br>135 (130, 140)<br>[126-142] | 136 ± 1.1<br>136 (134, 138.6)<br>[134-139] | 0.073 |
| **K<sup>+</sup> (mmol/L)** | 3.9 ± 0.3<br>3.8 (3.3, 4.5)<br>[3.1-5.1] | 3.7 ± 0.2<br>3.7 (3.2, 4.2)<br>[3.2-4.2] | < 0.000 |
| **Glucose (mmol/L)** | 7.2 ± 1.3<br>7.1 (5.1, 10.4)<br>[4.9-11.2] | 4.6 ± 0.6<br>4.5 (3.6, 6.4)<br>[3.6-6.7] | < 0.000 |
| **Lactate (mmol/L)** | 4.9 ± 1.6<br>4.7 (2.2, 8.7)<br>[1.2-9.6] | 1.2 ± 0.3<br>1.1 (0.7, 1.9)<br>[0.7-1.9] | < 0.000 |

pCO<sub>2</sub>: partial pressure of carbon dioxide, pO<sub>2</sub>: partial pressure of carbon dioxide, sO<sub>2</sub>: oxygen saturation, BE: base deficit, ctHb: total concentration of hemoglobin, ctBil: total concentration of bilirubin, Hctc: hematocrit

The table also illustrates the differences in maternal arterial blood gas parameters according to mode of delivery. Not surprisingly, significant differences were observed in most biochemical parameters between the groups. A significantly lower pH, pCO_2_, pO_2_, and sO_2_ were found in mothers giving birth vaginally as compared to planned CS. On the other hand, BD, total hemoglobin concentration (ctHb), hematrocrit (Hctc), total concentration of bilirubin (ctBil), potassium ion concentration (K^+^), glucose and lactate were significantly higher with VD as compared to planned CS. Comparison of nulli- and multipara women with VD showed significant differences in pO_2_, sodium ion concentration (Na^+^) and glucose levels (*P* < 0.02) (Table not shown).

Table 3 illustrates the influence of maternal characteristics and obstetrical interventions on acidbase parameters.

**Table 3.** Correlation of different maternal arterial acid-base values with respect to body mass index (BMI), smoking during pregnancy, the presence of hypertension/preeclampsia and the use of epidural anesthesia, oxytocin (intravenous) or nitrous oxide during labor.

|  |  |  | pH | P value | pCO<sub>2</sub> | P value | pO<sub>2</sub> | P value | ctHb (g/L) | P value | Glucose | P value | Lactate | P value |
| --- | --- | --- | --- | --- | --- | --- | --- | --- | --- | --- | --- | --- | --- | --- |
| **BMI** | **<18.5** | N=9 | 7.43 (0.05)<br>7.42 (7.36, -) | 0.41 | 3.31 (0.48)<br>3.53 (2.38, -) | 0.303 | 16.06 (8.85)<br>16.00 (15.10, -) | 0.145 | 141 (9)<br>141 (123, -) | 0.119 | 7.1 (2)<br>6.3 (5.5, -) | 0.47 | 4.4 (2)<br>3.4 (2.5, -) | 0.221 |
|  | **18.5-24.9** | N=128 | 7.42 (0.06)<br>7.41 (7.33, 7.54) |  | 3.29 (0.48)<br>3.36 (2.23, 4.21) |  | 15.63 (2.43)<br>15.6 (11.31, 21.97) |  | 135 (11)<br>137 (115, 156) |  | 7.2 (1.3)<br>7.1 (5, 10.4) |  | 5.0 (1.6)<br>4.8 (2.3, 8.9) |  |
|  | **25-29.9** | N=41 | 7.41 (0.06)<br>7.40 (7.32, 7.51) |  | 3.18 (0.61)<br>3.32 (1.08, 3.93) |  | 15.33 (1.99)<br>15.7 (10.36, 20.86) |  | 135 (9)<br>135 (118, 154) |  | 7.4 (1.3)<br>7.3 (5.1, 10.5) |  | 5.0 (1.9)<br>4.7 (1.2, 9.5) |  |
|  | **>30** | N=19 | 7.40 (0.04)<br>7.38 (7.34, 7.39) |  | 3.53 (0.48)<br>3.38 (2.79, -) |  | 14.71 (1.38)<br>14.8 (12.30, -) |  | 131 (11)<br>131 (108, -) |  | 7.0 (0.9)<br>7.0 (6.0, -) |  | 4.3 (0.9)<br>4.1 (3.2, -) |  |
| **Smoking** | **No** | N=173 | 7.41 (0.06)<br>7.41 (7.32, 7.54) | 0.398 | 3.3 (0.51)<br>3.37 (2.18, 4.22) | 0.805 | 15.46 (2.18)<br>15.60 (11.20, 20.48) | 0.949 | 135 (11)<br>135 (115, 156) | 0.926 | 7.26 (1.3)<br>7.1 (5.1, 10.4) | 0.731 | 4.9 (1.6)<br>4.7 (2.1, 8.4) | 0.421 |
|  | **Yes** | N=14 | 7.4 (0.05)<br>7.39 (7.34, -) |  | 3.37 (0.56)<br>3.22 (2.64, -) |  | 16.04 (3.36)<br>15.60 (12.6, -) |  | 134 (11)<br>138 (118, -) |  | 7.1 (0.9)<br>6.7 (5.9, -) |  | 5.4 (2)<br>4.9 (3, -) |  |
| **Hypertension/**<br>**Preeclampsia** | **No** | N=187 | 7.41 (0.06)<br>7.41 (7.32, 7.53) | 0.723 | 3.29 (0.52)<br>3.35 (2.19, 4.22) | 0.420 | 15.53 (2.29)<br>15.60 (11.60, 21.20) | 0.065 | 136 (11)<br>137 (116-155) | 0.063 | 7.2 (1.3)<br>7.1 (5.1-10.3) | 0.729 | 5.0 (1.7)<br>4.7 (2.1-8.7) | 0.998 |
|  | **Yes** | N=9 | 7.41 (0.04)<br>7.41 (7.33, -) |  | 3.40 (0.33)<br>3.51 (2.68, -) |  | 14.17 (1.09)<br>14.7 (10.30, -) |  | 130 (7)<br>130 (119, -) |  | 7.2 (1.7)<br>7.1 (5.7, -) |  | 4.9 (1.2)<br>5.2 (3.4, -) |  |
| **EDA** | **No** | N=159 | 7.41 (0.06)<br>7.40 (7.32, 7.53) | 0.047 | 3.29 (0.51)<br>3.34 (2.18, 4.23) | 0.872 | 15.72 (2.00)<br>15.70 (11.90, 19.55) | 0.01 | 136 (11)<br>137 (118, 155) | 0.407 | 7.2 (1.3)<br>7.1 (5.1, 10.5) | 0.977 | 5.0 (1.7)<br>4.8 (2.1-8.7) | 0.363 |
|  | **Yes** | N=55 | 7.42 (0.05)<br>7.42 (7.32, 7.54) |  | 3.28 (0.55)<br>3.39 (1.99, 4.22) |  | 14.90 (2.52)<br>15.10 (7.85, 21.67) |  | 135 (11)<br>135 (111, 157) |  | 7.2 (1.3)<br>7.1 (5, 10.4) |  | 4.7 (1.5)<br>4.5 (2.2, 8.7) |  |
| **NO<sub>2</sub>** | **No** | N=27 | 7.42 (0.07)<br>7.42 (7.32, -) | 0.291 | 3.23 (0.54)<br>3.13 (2.27, -) | 0.377 | 16.53 (3.01)<br>16.00 (11.70, -) | 0.061 | 133 (13)<br>135 (97, -) | 0.496 | 7.3 (1.3)<br>6.9 (5.8, -) | 0.836 | 4.9 (1.6)<br>4.3 (2.6, -) | 0.967 |
|  | **Yes** | N=177 | 7.41 (0.06)<br>7.41 (7.32, 7.53) |  | 3.30 (0.51)<br>3.35 (2.18, 4.22) |  | 15.42 (1.95)<br>15.60 (11.79, 19.46) |  | 136 (11)<br>137 (118, 156) |  | 7.2 (1.3)<br>7.1 (5.1, 10.4) |  | 4.9 (1.7)<br>4.7 (2.1-8.7) |  |
| **Oxytocin** | **No** | N=99 | 7.42 (0.05)<br>7.41 (7.33, 7.53) | 0.239 | 3.39 (0.50)<br>3.43 (2.25, 4.28) | 0.08 | 15.86 (2.17)<br>15.70 (12.03, 22.31) | 0.067 | 136 (12)<br>138 (112, 154) | 0.409 | 6.8 (1.2)<br>6.5 (5.0, 10.0) | 0.000 | 4.6 (1.5)<br>4.5 (2.1, 8.1) | 0.019 |
|  | **Yes** | N=115 | 7.41 (0.06)<br>7.41 (7.32, 7.55) |  | 3.20 (0.51)<br>3.28 (2.13, 4.14) |  | 15.20 (2.11)<br>15.40 (10.14, 21.05) |  | 135 (10)<br>136 (116, 158) |  | 7.6 (1.3)<br>7.4 (5.4, 10.6) |  | 5.2 (1.7)<br>4.9 (2.3, 8.8) |  |

“N” is the number of valid cases per parameter. For two group comparison Mann Whitney *U* test. For multiple group comparison Kruskal-Wallis test.
EDA: epidural anesthesia, NO<sub>2</sub>: nitrous dioxide, pCO<sub>2</sub>: partial pressure of carbon dioxide, pO<sub>2</sub>: partial pressure of carbon dioxide, ctHb: total concentration of hemoglobin

For biochemical parameters not shown in Table 3, the use of nitrous oxide gas (N_2_O) was found to significantly effect potassium (3.8 mmol/L versus 4.0 mmol/L, *P* < 0.000) and bilirubin levels (12.2 μmol/L versus 18.4 μmol/L, *P* < 0.012). Similarly, the use of oxytocin had a significant impact on sodium concentrations (134.6 mmol/L versus 135.9 mmol/L, *P* < 0.000) and oxygen saturation (98.1% versus 98.4%, (*P* < 0.01).

Calculated only for women with VD, multiple regression analysis with pH, pCO_2_, pO_2_, glucose and BD and lactate as dependent variables and the use of EDA, oxytocin and N_2_O respectively as independent variables, significant associations were found for the use of oxytocin and pCO_2_ (*P* = 0.016), glucose (*P* = 0.000) and lactate (*P* = 0.013) levels in maternal blood. Maternal arterial pH, pCO_2_ and lactate values correlated significantly to values in venous umbilical cord blood (*P* < 0.000) (Table not shown) with the correlation coefficient (R^2^) values as follows: pH R^2^ = 0.22, pCO_2_ R^2^ =0.07 and lactate R^2^ = 0.38. Although significant, the total duration of active “pushing” during the second stage of labor correlated poorly to both maternal and fetal lactate concentration (R^2^ =0.06, *P* < 0.000). In addition, we found no significant correlation between placental weight and fetal lactate concentration (R^2^ = 0.01, *P* = 0.431).

## Discussion

To the best of our knowledge, this is the first study to present reference values in maternal arterial blood at delivery. Although data collection and analysis were performed some years ago, the Radiometer ABL800 is still popularly used for blood gas analysis in many parts of the world making our results relevant today (13).

### Values in maternal arterial blood

Vaginal delivery is often compared to running an exhausting marathon for the delivering mother. To highlight the drastic effect on maternal arterial blood gas values during a vaginal delivery, we choose women undergoing planned CS as a control group. The significant differences in acid base values according to mode of delivery came as no surprise.

With VD, the sheer force of uterine contractions and bearing down results in impaired oxygenation and anaerobic glycolysis with lactate accumulation in both the mother and fetus (14, 15). The higher BD values in the VD group strengthened this observation, showing a tendency towards metabolic acidemia in the VD group. The lower pCO2 concentration can be explained by maternal hyperventilation, which is used as a compensatory mechanism for the metabolic component (respiratory compensated metabolic acidosis) to help keep maternal pH within a normal range. In addition, low maternal pCO_2_ facilitates the elimination of fetal pCO_2_ (Double Bohr effect) (15) thereby protecting the fetus from severe fetal respiratory acidosis. The lower hemoglobin concentration in the CS group is explained by the practice of giving a bolus of intravenous fluids when administering local anesthesia whilst higher glucose values in maternal arterial blood during vaginal deliveries can be attributed to a heightened sympathetic reaction prompting the body to release more glucose via glycogenolysis (16).

We were able to show significant differences in the VD group according to type of obstetric intervention used. EDA, inhibits neuronal feedback from sensory nerves in the uterus to the brain, resulting in a reduction in the endocrine pain response and decreasing adrenalin secretion from the adrenal medulla. There is also a reduction in oxytocin release from the pituitary gland which, together with decreased sympathetic response, results in maternal hypotension, motor blockade, low blood pressure and respiratory depression (7). Although we were unable to see any difference in pCO_2_ levels, pO_2_ levels were found to be significantly lower in women receiving EDA. We also saw higher pH and prominently lower lactate levels in the EDA subgroup, which can be explained by less effective muscle- including uterine contractions. Clinically, the use of EDA increased the likelihood of using synthetic oxytocin in order to augment uterus contractions and thus maintain normal “progress” during the first and second stages of labor (9). Stimulation with synthetic oxytocin, however, is not synonymous to normal physiological labor. Firstly, synthetic oxytocin does not cross the blood-brain barrier and secondly, during normal labor, oxytocin is released as small narrow peaks. Synthetic oxytocin is administered as a continuous infusion resulting in uniform levels which are suggested to influence the uterine muscle work with increased risk for hyperstimulation and lactate accumulation (17). The active transportation of the lactate ion into the maternal circulation leads to an increase in the level of lactate as seen from our results.

We explored the relationship of various maternal characteristics like BMI, smoking and hypertension on arterial acid base values and electrolytes. Although no significant changes could be shown, a larger study cohort than ours may be needed to investigate these characteristics further.

### Correlation between maternal and fetal lactate

We did not find a strong correlation between pushing time and maternal pH (not surprisingly since the value of pH is logarithmic) and lactate. In a smaller material. Nordström et al showed a sharp increase in both maternal and fetal lactate during the second stage of labor (18). The vast majority of fetal lactate is produced during this stage, and a prolonged stage of expulsion is associated with higher lactate concentrations in both umbilical cord- and fetal scalp blood (10, 20). Sub-analysis of our data showed that an increase in maternal lactate was associated with a corresponding increase in umbilical cord venous lactate values which, in turn, were positively associated with lactate values sampled from fetal scalp blood. However, the regression model only accounted for 38% of the venous fetal lactate increase, indicating that the rest must originate from the fetus itself. Transfer of lactate across the placenta may thus contribute minimally to the total fetal lactate concentration increase during the second stage of labor. The placenta is metabolic organ with lactate generation but we were unable to demonstrate any association between the weight of the placenta and fetal lactate concentration. Therefore, like previous studies, we concluded that the majority of fetal lactate was produced through endogenous lactate production within the fetus itself. This finding reinforces the use of fetal lactate from scalp blood sampling as a reliable tool in the evaluation of fetal distress (19, 20).

### Strengths and limitations

One of the major strengths of the current study was that blood sampling was conducted by trained professionals in both the groups. A specialist obstetrician (N.W) took fetal scalp lactate and maternal arterial blood gas samples in the VD group. In addition, all blood gas samples were analyzed immediately upon procurement, aided by a blood gas analyzer machine located within the Labor and Delivery department of the Hospital. Relevant obstetric and neonatal data was also entered directly after delivery into the study’s data base.

### Conclusions

Reference values for maternal arterial blood gases in vaginal deliveries for term pregnancies have been outlined with this study. We found that most arterial blood gas parameters varied significantly according to mode of delivery. In addition, different obstetrical interventions like the use of epidural anesthesia or synthetic oxytocin, resulted in significant changes in blood gas values. It may, therefore, be concluded that laboring women have altered biochemical parameters and reference values based on non-pregnant women should be interpreted with caution.

## Acknowledgments

Our sincere thanks to all the participating women in the study.

